# Mapping the sequence determinants of constitutive activity in GPCRs

**DOI:** 10.64898/2026.10.30.762644

**Authors:** Matthew K. Howard, Sara Whiteman, Patrick Rockefeller Grimes, Christian Macdonald, Aashish Manglik, Willow Coyote-Maestas

**Affiliations:** Department of Pharmaceutical Chemistry, University of California, San Francisco, San Francisco, CA, USA; Department of Bioengineering and Therapeutic Science, University of California, San Francisco, San Francisco, CA, USA; TETRAD graduate program, University of California, San Francisco, San Francisco, CA, USA; UC Berkeley-UCSF Graduate Program in Bioengineering, University of California, San Francisco, CA, USA; Department of Anesthesia and Perioperative Care, University of California, San Francisco, CA, USA; Chan Zuckerberg Biohub, San Francisco, CA, 94158, USA; Quantitative Biology Institute, University of California, San Francisco, San Francisco, CA, USA

## Abstract

G protein-coupled receptors (GPCRs) signal not only in response to ligands but also in their absence. This basal, or constitutive, activity has distinct physiological consequences and is the only functional measurement available for orphan receptors. It is also hard to compare receptors because each is reported relative to its own baseline. Here, we measure basal activity on a single scale across the human GPCR family. Pairing a barcoded library of all human GPCRs with a transcriptional cAMP reporter, we scored 901 receptor constructs in a pooled experiment. Basal activity forms a continuous distribution, and many of the highest-scoring receptors have prior reports of ligand-independent activity, with orphans being strongly represented. We then exhaustively mutated one of them, GPR161, and measured its effects on protein abundance, surface expression, and cAMP signaling. Of the 528 positions, only 10 specifically alter signaling independent of expression. Constraints on basal signaling are dominated by receptor biogenesis, which requires multiphenotype scanning to resolve. Together, these datasets place basal activity on a common scale across the GPCR family and provide a template for dissecting it at residue resolution.

## Introduction

G protein-coupled receptors (GPCRs) are the largest family of cell surface receptors in the human genome. They detect a chemically diverse set of extracellular stimuli and transduce them across the membrane by coupling to heterotrimeric G proteins and arrestins, which in turn regulate second-messenger levels and downstream programs ^1^. Conserved across all GPCRs is a heptahelical fold whose rearrangement converts stimulus detection into transducer coupling. The prevailing model of ligand activation of receptors is that ligands bias the conformational ensemble toward states that can productively signal. It follows that receptors populate signaling-competent states at some frequency even in the absence of a stimulus, and for many receptors this is sufficient to produce measurable signaling. This basal, or constitutive, activity is the baseline that agonists raise and inverse agonists lower, and it can have distinct cellular and physiologic consequences^1–3^. For instance, mutations that raise basal activity can cause a variety of developmental and endocrine disorders, and variants that lower it are associated with obesity and cancer predisposition, among other conditions^4,5^. For orphan receptors, those that have no known ligand, basal output is the only functional measurement available.

Basal activity is challenging to compare between receptors, as it is often only visible as a difference between two conditions. One example is comparing a non-expressing cell to a receptor-expressing cell, or examining the drop in signal produced by an inverse agonist. These quantities are also measured in a wide variety of assays (e.g., cAMP accumulation, reporter transcription, GTPγS binding, or transducer recruitment), which are difficult to compare directly. Many receptors have been studied, but we lack comprehensive panels of basal activity across the entire GPCR family. Assays have been developed to measure many receptors at once, which has changed the questions that can be asked of the family.

Several platforms have mapped beta-arrestin recruitment across the druggable GPCRome and transducer coupling across several hundred receptors^6,7^. Previous work has begun to develop genetically barcoded receptor libraries to enable sequencing-based readouts of receptor activity^8^, and our group has recently developed a fully pooled method to measure expression differences across all human GPCRs^9^. In all accounts, there are no GPCRome-wide measures of basal activity.

While a GPCRome-scale measurement can identify which receptors have basal activity, it does not identify which residues govern basal receptor activation. An ideal approach would be to screen all possible receptor mutations for their effects on receptor activity. Deep mutational scanning (DMS) enables pooled measurements of protein function across a comprehensive mutational library^10^. DMS has been applied to several ligand-regulated GPCRs to study activation mechanisms^11–15^. However, receptor variants may alter multiple receptor phenotypes upstream of receptor activation, thereby confounding the results^16^. To address this, a multiphenotype screen can be used to assess effects on both expression and signaling, thereby identifying variants and the mechanism(s) by which they alter receptor function. Comparing mutational effects on constitutive activity and expression could reveal how variants govern basal receptor signaling.

Here we approach this question on two fronts. We develop a pooled, barcoded GPCR library with a transcriptional cAMP reporter and use it to measure basal activity across the entire human GPCRome. The screen reports relative cAMP activity, enabling, for the first time, a measure of basal activity across the entire family. Among the top basally-activating receptors are many with previously reported ligand-independent activity, including orphan receptors. GPR161 ranks highly due to its constitutive activity, offering a unique opportunity to study basal signaling at the residue level using DMS, especially given its well-characterized role in development and existing structural work identifying features that may govern its activity^17–19^. We therefore measured variant effects on GPR161 across several phenotypes to determine effects on expression and signaling.

## Results

### Pooled measurement of basal cAMP signaling across the GPCRome

Many GPCRs are active or partially active in the absence of a stimulus. This basal, or constitutive, activity is an important feature in certain biological and physiological contexts and is often targeted for pharmacological modulation^3,4^. Basal activity is also hard to compare across receptors, as it is typically reported relative to each receptor’s own baseline in a particular assay, so the results from one experiment cannot necessarily be compared to another. Here, we provide a first step towards measuring basal activity across the human GPCR family.

Measuring basal activity across a receptor family requires a library of all receptors and a signaling readout that facilitates their comparison. We use a recently developed GPCR library from Tedman et al.^9^, which contains the sequences of all human GPCRs and many splice isoforms thereof in a single plasmid pool (946 constructs in total). Each receptor in the library is paired with a single unique 10-nucleotide barcode and attB site, enabling integration into cell lines. Crucially, the vectors contain restriction sites that enable subcloning of the entire receptor library into different backbones. We use these sites to transfer the GPCR library into a vector containing a transcriptional cAMP reporter via Golden Gate assembly (**Figure 1a**). This reporter enables the accumulation of an eGFP fluorescent signal in response to cAMP accumulation and has been used in previous pooled assays of GPCR activity, e.g., deep mutational scanning ^15^. Combining the GPCR library with this reporter enables the measurement of cAMP activity across the entire GPCR family.

**Figure 1:**
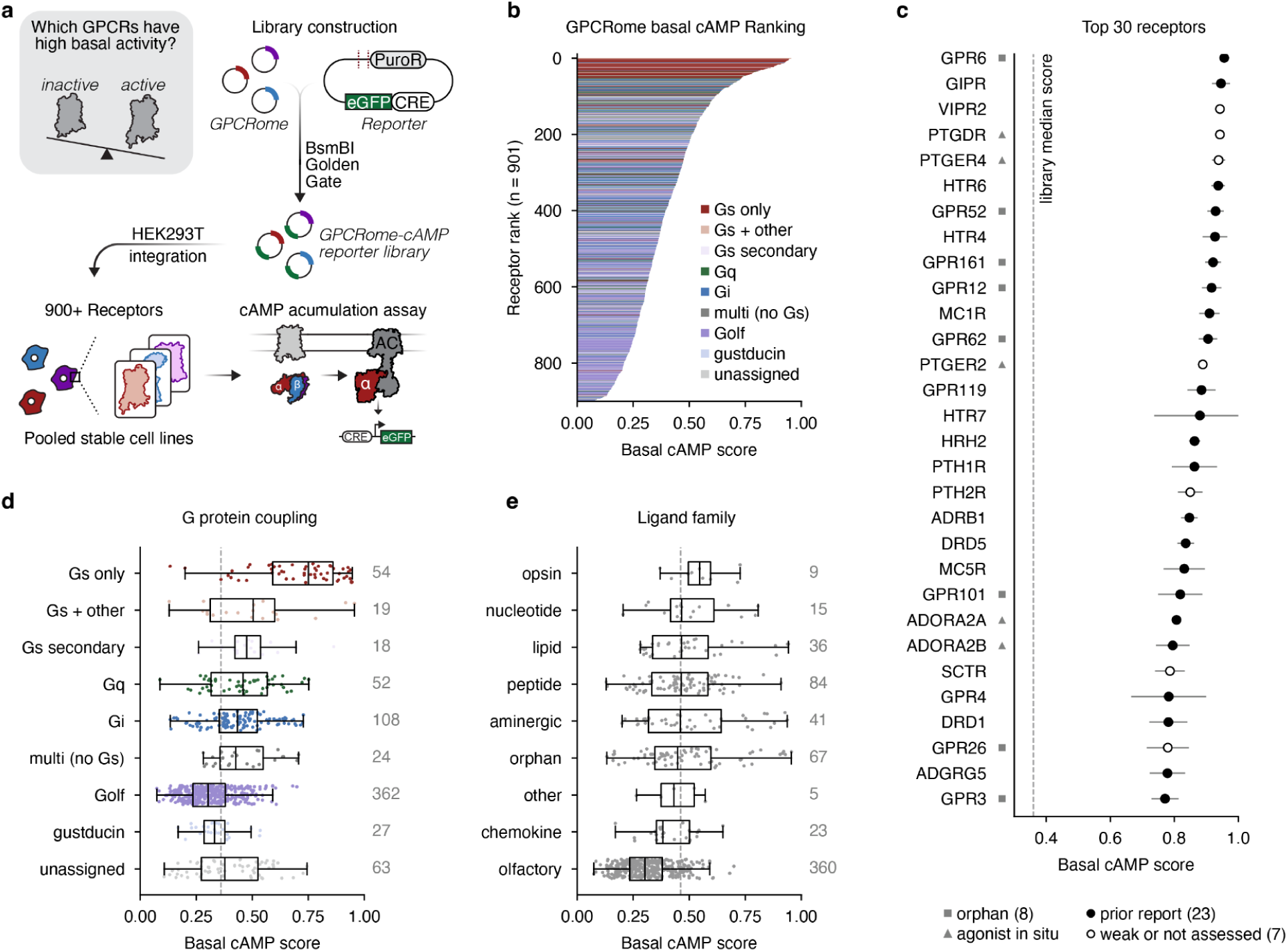
Basal cAMP signaling across the GPCRome. (a) Platform to capture ligand-independent cAMP signaling across the human GPCRome. The GPCRome receptor library from Tedman et al.^9^ was cloned into a fluorescent cAMP reporter plasmid previously used for Gs-coupled receptor mutational scanning studies^15^. (b) Relative cAMP signaling across the GPCRome. Scores are scaled from 0 to 1, where higher values indicate greater cAMP reporter activation. Receptors are colored by G protein coupling. (c) Top 30 receptors from **b**. Bars represent standard deviations. Orphan receptors and receptors with potential *in situ* ligand confounds are indicated. Receptors with previously reported constitutive activity are indicated, to the best of our curation abilities. (d) Box plots showing the distribution of GPCRs across each G protein class. (e) Box plots showing distribution when class A GPCRs are grouped by ligand class.

The resulting plasmid library was integrated as single-copy genomic insertions into HEK293T cells via a Bxb1 landing pad system, ensuring only one receptor construct is delivered per cell (**Figure 1a**)^20^. Expression of the receptor library is under doxycycline control. To resolve differences in basal cAMP signaling across the library, we induced expression for 2 days and accumulated eGFP/cAMP signals for 24 hours (see methods) prior to assaying by fluorescence-activated cell sorting (FACS) (**Supplemental Figure 1**). Receptors are assigned a quantitative score representing their inferred distribution across sort bins, based on barcode counts from sequencing (**Supplemental Figure 1b-e**). Across replicates, we observe a strong correlation (r = 0.90) and recover reliable measurements for 901 distinct receptors (**Supplemental Figure 1b-e**).

### Determinants of basal cAMP activity across the GPCR family

Several features of the resulting activity profiles provide confidence that the screen faithfully reads constitutive Gs/cAMP signaling across the GPCRome library. As expected for a Gs/cAMP reporter assay, we observe that Gs-coupled receptors score higher than receptors coupled to other transducers (e.g., Gq, Gq, etc)(**Figure 1b-d**)^21^. All of the top 30 scoring receptors are Gs-coupled. Of these top 30 receptors, 23 have previous reports of ligand-independent activity (**Figure 1c**). Gs-coupled receptors are not compressed toward the upper extreme; we observe them across the full range of cAMP levels (**Figure 1d**). Beyond Gs-coupled receptors, we observe that Gi, Gq, and other couplings display a range of scores, but sit closer to the library median, with none reaching as high as the top Gs receptors (**Figure 1d**). Finally, we note that olfactory receptors (360 of the 727 ranked canonical constructs) couple to Golf (but couple to Gs in HEK293T systems) and display the lowest scores, likely because they express poorly and thus do not generate as much cAMP as more well-expressing GPCRs.

We next asked whether receptor families differ in their propensity to generate basal cAMP. Interestingly, opsins have the highest median cAMP signal, but several other ligand families contain individual receptors with much higher basal cAMP signaling levels (**Figure 1e**). Lipid, peptide, aminergic, and orphan receptor families each contain receptors that exhibit high cAMP signaling (**Figure 1e**). High basal cAMP activity is therefore not a unique property of one receptor family.

Several of these hits are potentially confounded by factors that a pooled cAMP screen cannot resolve. A few class B receptors, such as GIPR (rank 2), VIPR2 (3), PTHR1/2R (17/18), and SCTR (25), all rank highly, and agonist-independent cAMP is known to scale with receptor expression ^22,23^. A second set of receptors may be activated by endogenous agonists produced *in situ*. HEK293 cells can express prostaglandin^24^, which may drive activation of three prostanoid receptors in the top 30: PTGDR (4), PTGER4 (5), and PTGER2 (13). Additionally, the adenosine receptors ADORA2A (23) and ADORA2B (24) face the same confound. HEK293 cells also produce adenosine, which may be driving adenosine receptor activation^25,26^. GPR4 (26) is a proton-sensing receptor rather than a constitutively active receptor, and its pH_50_ for cAMP is close to 8, so at the pH at which this experiment was run (∼7.4), it is activated^27^. GPR119 (14) responds to oleoylethanolamide and to lysophosphatidylcholine^28,29^, either of which the serum in the medium may supply. These findings, therefore, do not indicate inaccurate measurements but rather reflect idiosyncrasies of cell-based assays in general and the challenge of performing pooled assays in cells that produce these ligands.

Several receptors with distinct profiles remain after accounting for potential confounds. In particular, well-characterized orphans are strongly represented at the top. GPR6 ranks highest, along with its family members GPR12 (10) and GPR3 (30). This family signals through Gs at levels comparable to a fully activated receptor with no ligand added^30,31^. GPR52 (7) follows, and its mechanism is structurally characterized. GPR52’s extracellular loop 2 (ECL2) folds into the orthosteric pocket and acts as a tethered agonist^32^. GPR62 (12) lacks a canonical DRY motif, which has been implicated in its constitutive cAMP signaling ^33^. GPR101, whose duplication causes X-linked acrogigantism, is also constitutively Gs-coupled^34,35^. GPR161 ranks 9th. GPR161 constitutively signals through Gs, raising cAMP to repress Gli transcription factors in Hedgehog signaling^17^, and its output is controlled by removing it from the primary cilium^36^.

Isoforms of several receptors exhibit differences in basal cAMP levels (**Supplemental Figure 2**). Two cases occur among receptors with high basal cAMP signaling: GPR161 and ADORA2A (**Supplemental Figure 2a**). GPR161 is represented by its canonical transcript and four alternative isoforms, and ADORA2A by its canonical transcript and five isoforms. In each receptor, one alternative isoform preserves all seven predicted transmembrane helices: for GPR161, this is ENST00000537209, which carries a 20-residue N-terminal extension, and for ADORA2A, it is ENST00000444262, a C-terminally truncated form. Both retain full basal cAMP output but differ in relative surface expression (**Supplemental Figure 2b-d**). The GPR161 isoform shows a reduction in surface expression relative to the canonical form, while the ADORA2A isoform is expressed at canonical levels. The remaining isoforms of both receptors lose three or more of the seven helices and fall near the library median with sharply reduced surface expression (**Supplemental Figure 2b-d**). Other receptors show comparable distributions across their isoforms, suggesting this platform can be used to ask how natural sequence variation within a gene affects expression and activity, not only how receptors differ from one another.

In conclusion, the GPCRome basal cAMP screen identifies known biology and nominates receptors that have high basal cAMP in HEK293T cells. While not necessarily a reflection of how these receptors might behave in a physiologic context, this assay provides a platform to study factors that regulate the activity of these candidate receptors in the absence of ligand.

### Deep mutational scan of GPR161 expression and signaling

The GPCRome library is a resource for comparing distinct receptor sequences and activity levels. However, we were curious to achieve a granular, residue-resolved understanding of how sequence sculpts receptor activity in an unliganded system. For this purpose, we chose a single receptor and extensively mutated it to understand how individual variants contribute to its function. GPR161 was a natural choice for this experiment. GPR161’s constitutive activity is the functional property that matters for its role in development, where ciliary GPR161 holds the Hedgehog pathway off until Smoothened drives the receptor out of the cilium^17,36^. Germline loss-of-function predisposes to infant Hedgehog group medulloblastoma ^37^, and rarer variants have been reported in pituitary stalk interruption syndrome ^38^ and spina bifida ^39^. GPR161 is deeply conserved across vertebrates^19^, and in our GPCRome screen, it is amongst the highest-signaling receptors (**Figure 1c**).

We constructed a variant library of GPR161 using the DIMPLE pipeline^40,41^. At each of the 528 positions covered by the library (positions 2-529), we encoded all missense mutations, a synonymous variant, and a single amino acid deletion. Similar to the GPCRome library, the GPR161 mutational library was integrated into a cAMP reporter vector and delivered as single-copy insertions into HEK293T cells via a Bxb1 landing pad system^15,20^. We observed 99.1% of the designed variants in the final cell line library (**Figure 2a, Supplemental Figure 3**).

**Figure 2:**
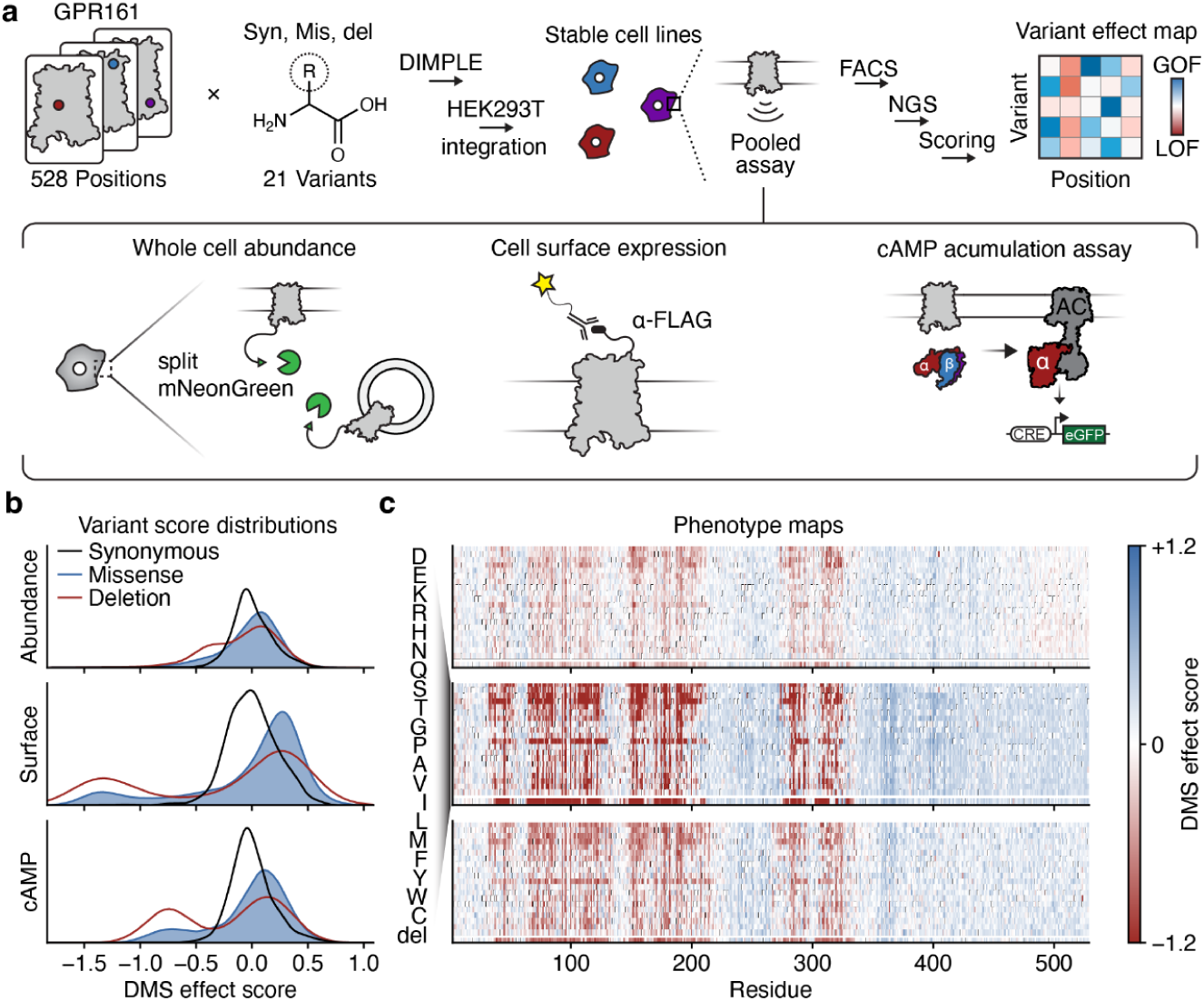
GPR161 DMS overview. (a) Deep mutational scanning workflow for GPR16 library design, cloning, cell line integration, and screening. The GPR161 variant library was screened across 3 phenotypes: cell surface expression, whole-cell abundance, and cAMP signaling. (b) Density distributions of variant effects across each phenotypic assay. Scores for synonymous variants are centered at zero. Positive scores indicate increases in fitness for a given assay. Negative scores represent loss of fitness. (c) Heatmaps of individual variant effects across each phenotypic assay. Positive scores indicate increases in fitness for a given assay. Negative scores represent loss of fitness.

Variant effects were assessed across three phenotypic assays that monitor whole-cell protein abundance, cell-surface expression, and cAMP signaling (**Figure 2a-b, Supplemental Figure 3**). Abundance was monitored using a split mNeonGreen complementation assay, similar to previous VAMP-seq approaches applied to soluble and membrane proteins^42,43^. The C-terminus of the GPR161 library was tagged with mNeonGreen11, the 11th strand of the beta barrel. Transient transfection of mNeonGreen1-10 yields a fluorescent signal for variants that are expressed at any location in the cell, whereas misfolding variants result in diminished mNeonGreen signal. Cell surface expression was monitored using antibody staining of an N-terminal FLAG tag. Comparing abundance with surface expression enables the identification of residues that constrain or potentiate surface expression. Finally, cAMP production was measured using the same assay employed for the basal cAMP GPCRome screen. All assays followed a similar FACS selection scheme (see methods; **Supplemental Figure 3**). Effect scores for each assay were calculated from variant counts derived from NGS using the Lilace algorithm^44^. Scores for each phenotype are centered at zero, and negative scores indicate loss of function and positive scores indicate an increase in function relative to the synonymous mean. Read together, these three assays allow attribution of a mechanism explaining why a variant might behave differently from the wild-type receptor.

Across the three mutational scans, we achieve good resolution between functional and non-functional variants. This is highlighted by the deleterious effects of the missense and deletion variants in the transmembrane domain, while the intervening loops and n/c-termini remain relatively unconstrained in each assay (**Figure 2b**).

### Decomposing variant effects across three phenotypes

The three measurements for variant effects on GPR161 abundance, surface expression, and cAMP signaling are correlated, but non-redundant. Across the 9,927 missense variant scores in all three assays, surface expression track abundance at ρ = 0.63, and cAMP at ρ = 0.66 (**Figure 3a-b**). To understand which variants have effects on a given phenotype, we define a loss-of-function cutoff for each that corresponds to two standard deviations of the synonymous variant population (**Figure 3a-b**).

**Figure 3:**
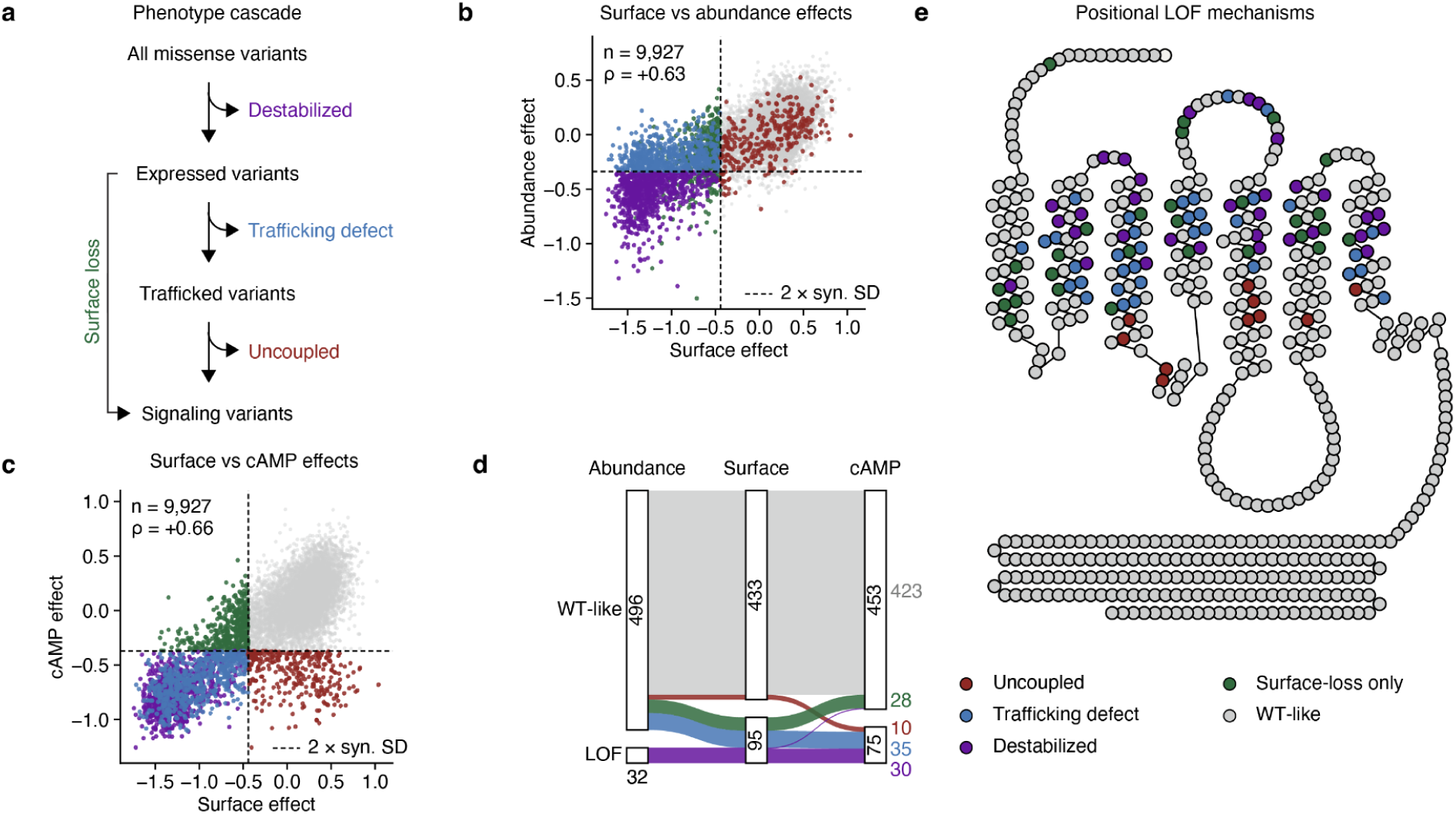
GPR161 variant effects across phenotypes. (a) Schematic for characterizing how mutations alter GPR161 based on DMS effects (b) Comparison of variant effects on GPR161 protein abundance and cell surface expression. (c) Comparison of variant effects on GPR161 surface expression and cAMP signaling. (d) Sankey plot demonstrating how variants alter each assayed DMS phenotype. (e) GPCR snake plot showing how positions are classified based on **a**.

Of the 10,927 variants scored in all three assays, 1,880 (17.2%) lose basal cAMP signaling. The majority of these are not signaling-specific mutants, as many also exhibit expression defects. For 889 variants, all three phenotypes have low scores, meaning the receptor is not made and therefore cannot signal. For 654, abundance remains intact, while surface expression and cAMP are lost, implying that the receptor is made in cells but not delivered to the cell surface. S88P and W182G are examples of the second class that have wt-like abundance but fail to traffic to the cell surface. A further 452 variants lose surface expression while leaving cAMP intact, which implies that diminished surface expression does not necessarily abolish measurable signaling. After accounting for effects on expression and abundance, 309 variants specifically lose basal cAMP signaling (2.8% of the total library).

At the position level, we averaged missense effects and applied the same 2-standard-deviation cutoff to the per-position mean. Of the 528 positions, 423 are wild-type-like in all three assays; 32 lose abundance, 35 lose surface expression and cAMP while retaining abundance, 28 lose surface expression alone, and 10 lose cAMP signaling with expression intact (**Figure 3c-d**). In an expression-matched system, there may be more positions or variants with specific effects on cAMP signaling. However, the effects on expression percolate through our mutational scanning phenotypes, masking them. Constraint on basal signaling in GPR161 appears overwhelmingly due to making and delivering the receptor, and only ten positions in the entire protein are constrained specifically for coupling itself.

### Structural determinants of GPR161 expression and signaling

We were curious to provide a structural rationale for the positions identified as having large effects on GPR161 expression and signaling. We mapped these positions onto the active-state GRP161-Gs structure^19^, which resolves 268 of the 529 assayed positions.

Looking first at expression, we observe large defects in expression throughout the transmembrane domain. Specifically, transmembrane helices 2, 3, and 4 have the greatest effects on expression when mutated (**Figure 4a-b**). Interestingly, ECL2 carries a similarly large effect on expression when mutated (**Figure 4a-b**). Perhaps this loop helps stabilize the receptor when it is folded into the orthosteric site, beyond its role as a tethered internal agonist.

**Figure 4:**
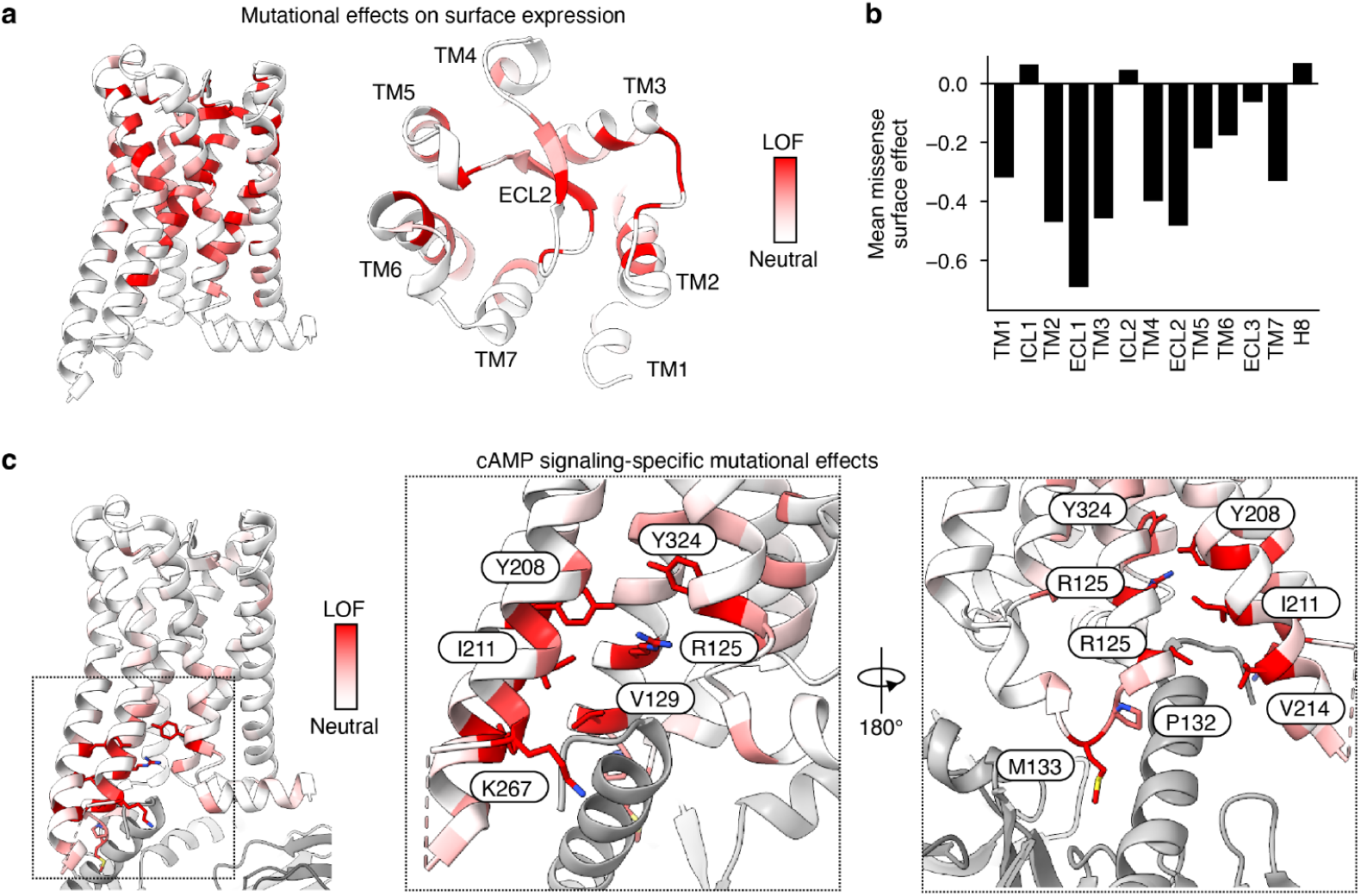
Structural interpretation of GPR161 variant effects. (a) Mapping of per-position variant effects on surface expression mapped onto the GPR161 structure (PDB: 8SMV). (b) Variant effects on surface expression by receptor region. (c) Mapping of per-position cAMP-specific effects mapped onto the GPR161 structure (PDB: 8SMV).

**Figure 5:**
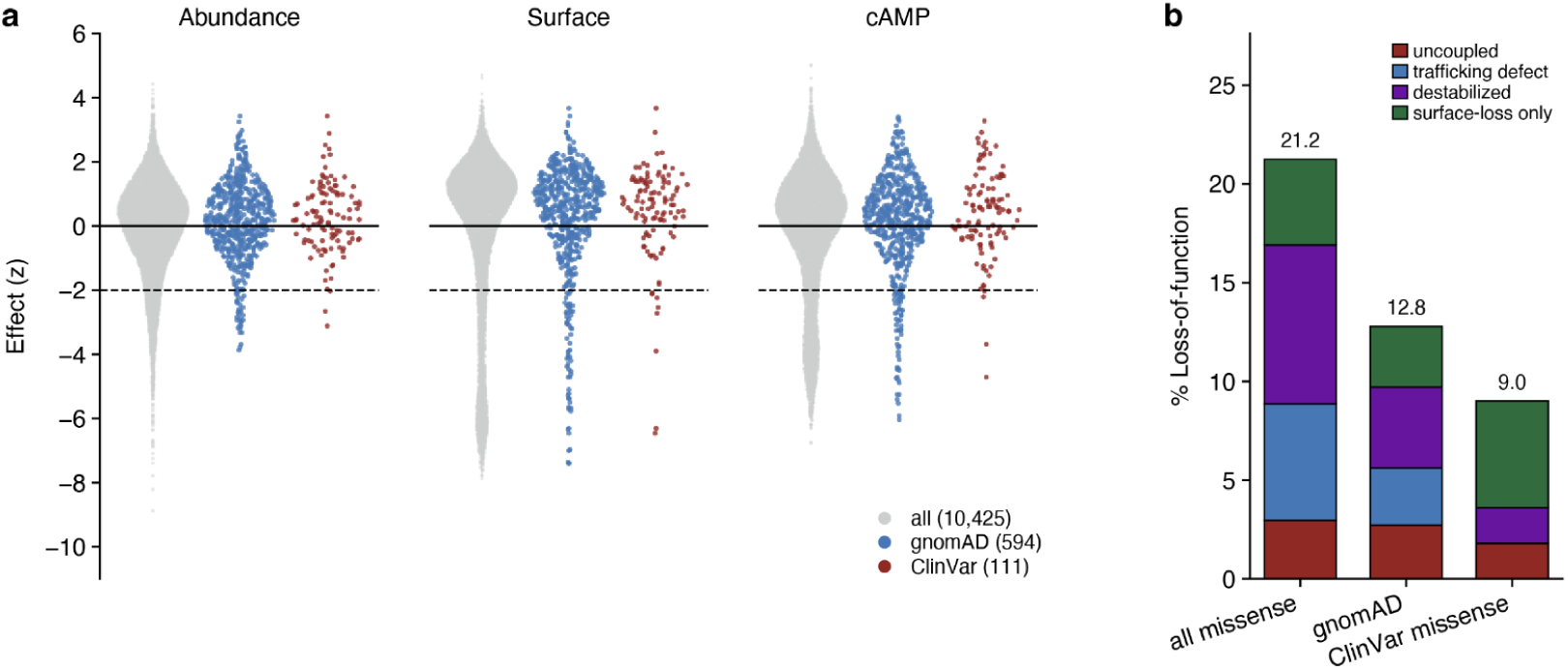
Interpretation of human GPR161 variants. (a) Mutational scanning identifies phenotypes altered by human variants present in gnomAD and ClinVar databases. Variants below the −2 SD dashed line are considered loss-of-function in each assay. (b) Mechanistic interpretation of GPR161 variants present in gnomAD and Clinvar.

Signaling-specific effects are concentrated in the G protein binding site (**Figure 4c**). Ten positions are contained in this set: R125^3.50^, V129^3.54^, P132^ICL2^, M133^ICL2^, Y208^5.58^, I211^5.61^, V214^5.64^, A215^5.65^, K267^6.32^, and Y324^7.53^. Superscripts denote the Ballesteros-Weinstein/GPCRdb common numbering system for class A GPCRs^45,46^. This set contains the DRY motif arginine, the tyrosine that packs against it in the active state, two positions in ICL2, and the NPxxY tyrosine, with nothing scattered elsewhere in the protein. The scan thus recovers the core class A GPCR activation machinery.

A prior structural study identifies two motifs involved in constitutive activity: ECL2, which occupies the canonical orthosteric site, and an extrahelical cholesterol adjacent to TM6 and TM7^19^. In the mutational scanning data, neither location shows a cAMP-specific effect. Rather, we see that they have large effects on receptor expression, so in this assay, the cAMP loss is likely due to reduced expression. It remains possible that these motifs are involved in activation, but because they are essential for expression, it is challenging to distinguish this. Perhaps an expression-matched experiment could resolve this disagreement.

Similarly, the extrahelical cholesterol site contains no expression-independent cAMP LOF positions. The Site is relatively small, so power may be limited, but across a saturating number of mutations, we find that W327 tolerates all 19 substitutions with no loss of basal cAMP or surface display. The previously reported single mutants at this site are indistinguishable from WT in the cAMP assay, and the 323A/327A/332A triple mutant cannot be directly compared to the single-variant DMS results, so we cannot determine whether any epistatic interaction underlies the reported phenotype. Additionally, any ciliary-dependent functional effects are not captured in these assays, which were done in non-ciliated HEK293 cells.

### Human variation tracks the mutational scan

We next asked how our variant effect maps relate to the genetic variation that exists in humans. Of the 602 missense alleles in gnomAD^47^, 593 encode unique proteins, and 587 of these are scored in our assays. ClinVar^48^ contributes an additional 112 missense variants, and we recover mutational scores for 111 of them. In total, we measured variant effects for 603 human alleles in the mutational scan.

Damaging variants are depleted among the observed human alleles. Across the missense variants in the library, 16.9% are loss-of-function in the cAMP assay and 18.3% in the surface expression assay. Among the gnomAD-observed set, those fractions are 9.5% and 10.0%; among singletons, 11.4% and 12.5%; and among alleles seen more than ten times, 2.2% and 5.5%. ClinVar missense variants, which are dominated by variants of uncertain significance rather than by established pathogenic alleles, account for 3.6% and 7.2%, respectively. This suggests that our mutational scan can separate alleles that persist in the human population from those that do not.

Fourteen observed human variants carry a cAMP-specific loss-of-function phenotype. Two of these, V129L and I211N, occur in the core of the receptor. Neither variant is annotated in ClinVar. Two ClinVar variants of uncertain significance (VUS), V80A and S316I, carry a large loss of function in the expression assays. Interestingly, two variants with a medulloblastoma annotation, T51N and V228F, are wt-like across all assays. Overall, our screen can stratify human variants and assign mechanisms to them.

## Discussion

These experiments provide complementary views on basal signaling in GPCRs. Measuring the activity of over 900 GPCRs together reveals how activity levels compare between receptors and identify receptors with unique signal profiles. We observe that basal activity forms a continuous distribution in which most receptors show little activity, while a few show large amounts. In part, this is due to measuring only one of the diverse signaling pathways that can be activated by a GPCR. Interestingly, when looking at Gs-coupled receptors here and the overall library, there is little correlation between receptor expression and basal signaling. This reflects differences among receptors and is not in conflict with the expression-level dependence observed in individual receptors. Many receptors at the top of the distribution are likely reading out unavoidable ligand-driven activity by ligands in cell culture media and/or produced by the cells themselves. Orphan and constitutively active receptors are among the cleanest hits, as they are active in the absence of a ligand.

In our forays with the orphan receptor GPR161, we find that mutations primarily drive differences in expression rather than directly modulating signaling output. Of the 528 positions assayed in the mutational scan, only 10 have effects that are specific to cAMP signaling. These positions cluster in and around the interface with the G protein, the surface that transmits activation in GPCRs. While unsurprising that these residues light up similarly in GPR161 relative to other class A GPCRs that have been subject to DMS, it is interesting that regions like ECL2 have little direct influence on cAMP that doesn’t result from decreases in receptor expression^11–15^. In part, this demonstrates the utility of performing mutational scanning across multiple phenotypes, especially in the absence of distinct stimuli, to discern residues involved in activation and to determine how these mechanisms may inform disease processes.

Basal activity is often treated as a baseline to subtract rather than as a property to measure. However, it provides a window into the balance between the receptor’s inactive and active ensembles. In an agonist-driven system, a mutation that reduces signaling may weaken ligand binding or impair activation; in the absence of ligand, that ambiguity disappears. DMS provides a unique opportunity to see how this property is tuned, provided effects are read along more than one phenotype. This is particularly true for GPR161, where we find receptor folding and trafficking can mask constraints on activation. Applied across a set of receptors, maps of this kind will separate the residues that set the activation equilibrium in general from those particular to a single receptor’s fold.

These studies provide a step towards understanding the sequence-level underpinnings of basal activity in GPCRs. Looking forward, these approaches are readily extendable for exploration in fundamental GPCR biophysics and discovery biology. In this work, we assay GPCR activity along a single vector, Gs/cAMP signaling. This ignores the pluridimensionality of GPCR signaling activity. We envision that similar approaches may be used to assay along other pathways. Platforms for pooled screening of GPCRs have already been developed for Gi, Gq, and arrestin recruitment^11–13^. In time, further development may occur to enable measurement of intracellular localization, trafficking, and signaling *en masse*. Cellular background also poses an area for exploration. Assaying different parameters in context-specific cell lines may prove fruitful. For instance, olfactory receptors may require factors uniquely present in olfactory sensory neurons, and other ciliary receptors would be better suited for study in NIH3T3 or other model ciliated cell lines. Regulators of these phenotypes may also be uncovered by coupling these readouts to additional perturbations, such as CRISPR. As technology and assay platforms such as these develop, there remains fertile ground to further interrogate the complexities of GPCR biology.

## Acknowledgments

We thank members of the Coyote-Maestas and Manglik labs for their helpful feedback and discussion as we conducted this work. We acknowledge the Astera Institute Hidden Science Competition for motivating us to release these datasets. We acknowledge funding support from the following sources. National Institutes of Health, National Institute of General Medical Sciences 5T32GM139786 (MKH), NIH National Institute on Drug Abuse DA056354 (WCM), Chan Zuckerberg Biohub (WCM and AM), Howard Hughes Medical Institute Hanna Gray Fellowship (WCM), Shurl and Kay Curci Foundation (WCM), Hypothesis Fund (WCM).

## Author contributions

MKH, WCM, and AM conceived the project. MKH, PRG, and WCM generated and cloned the GPR161 deep mutational library. MKH cloned the GPCRome library and generated all GPCR and GPR161 DMS cell lines. MKH and SW performed GPCRome basal signaling experiments. MKH designed and performed deep mutational scan experiments with assistance from PRG. MKH and CBM processed raw next-generation sequencing data. MKH analyzed datasets with input from WCM and AM. MKH prepared figures. MKH wrote the manuscript, with edits and approval from all authors. WCM and AM supervised the overall project.

## Data availability

- GPR161 variant count data, DMS scores, and scripts used to analyze data and prepare figures have been deposited on Github:https://github.com/matthewkarlhoward/2026_GPR161_mutational_scan and Zenodo: https://doi.org/10.5281/zenodo.23682577.
- GPCRome receptor count data, scores, and scripts used to analyze the data and prepare figures have been deposited on Github: https://github.com/matthewkarlhoward/2026_GPCRome_basal_cAMP and Zenodo: https://doi.org/10.5281/zenodo.23682602.
- GPCRome basal signaling scores are included in **Supplemental Table 1**.
- DMS scores are included in **Supplemental Table 3** and deposited on MaveDB: urn:mavedb:00001323-a-1, urn:mavedb:00001323-b-1, urn:mavedb:00001323-c-1.
- Raw sequencing reads for the GPCRome basal cAMP screen have been deposited in the NCBI Sequence Read Archive under BioProject accession PRJNA1551037.
- Raw sequencing reads for the GPR161 mutational scan have been deposited in the NCBI Sequence Read Archive under BioProject accession PRJNA1551518.

## Methods

### GPCRome library construct design and cloning

A pooled plasmid library containing human GPCR sequences and isoform variants was a gift from Jonathan Schlebach (Purdue University)^9^. In this library, each GPCR is constructed as follows: [BsmBI]-[UMI]-[attB]-[GPCR]-[BsmBI] inside the backbone. Here, we use Golden Gate assembly (BsmBI) to transfer the [UMI]-[attB]-[GPCR] cassette into a modified vector containing an [IRES]-[PuroR] downstream of the GPCR and a cAMP reporter system that has been previously used for GPCR mutational scanning experiments^15^. Construct sequences are included in **Supplemental Table 1**.

### GPR161 deep mutational library generation

The DIMPLE platform was used to generate the GPR161 deep mutational library^40,41^. Briefly, we designed the library to contain all missense mutations at each position in GPR161. We additionally included synonymous mutations and single-amino-acid deletions at each position. These mutations were encoded in oligos with flanking BsaI sites and then ordered as a SurePrint Oligonucleotide library (Agilent Technologies) (**Supplemental Table 2**). DNA was resuspended, and the sublibrary fragments were amplified using PrimeStar GXL DNA polymerase and fragment-specific primers (**Supplemental Table 2**. These reactions underwent PCR cleanup using Zymo Clean and Concentrate-5 kits. The cDNA sequence of GPR161 WT was synthesized by Twist Bioscience in their High Copy Number Kanamycin backbone; BsmBI and BsaI cut sites were removed via silent mutations. For each library fragment, this plasmid was amplified to add BsaI sites, gel-purified, and the corresponding oligo sublibrary was assembled using BsaI-mediated Golden Gate assembly. These reactions were cleaned using and electroporated into MegaX DH10B cells, recovered for 1 hour, and added to 30 mL LB + Kanamycin and grown while shaking at 37 C until they reached OD 0.6-0.7. DNA was isolated using a Zymo Zyppy Plasmid Miniprep kit. Each sublibrary was quantified using Invitrogen Qubit dsDNA HS assay kit and pooled in equimolar ratios. This pooled library was then assembled into a BxBI landing pad-compatible cAMP reporter vector containing a GSGSGS-P2A-PuroR cassette for positive selection and a CRE-GFP reporter ^15^.

### Stable cell line library generation

The HEK 293T LLP-iCasp9 cells used in this study were a gift from Doug Fowler (UW)^20^. Cell lines for GPCRome and GPR161 DMS libraries were generated as follows. 1ug of library DNA was cotransfected with 1ug BxB1 recombinase (pCAG-NLS-BxB1, Addgene #51271) using 3.75uL lipofectamine 3000 and 5uL P3000 reagent in each well of a 6 well plate. For the GPR161 DMS library, 18 wells were transfected in parallel, and for the GPCRome, 6 wells were transfected. Cells were cultured in “D10” media (DMEM, 10% dialyzed FBS, 1% sodium pyruvate, and 1% penicillin/streptomycin) inside humidified incubators at 37 °C and 5% CO_2_. The landing pad in the cell line contains a Tet-on promoter upstream of the BxB1 recombination site and a split rapamycin analog inducible dimerizable Casp-9. Two days after transfection, we induce with doxycycline hyclate (2 µg/mL) and treat with 10 nM AP1903. Recombined cells have shifted the iCasp-9 cassette out of frame, while unrecombined cells will express the cassette and, upon treatment with AP1903, die from iCasp-9-induced apoptosis. Cells were selected for 2 days in AP1903, after which they were transitioned back to D10 supplemented with doxycycline. After two days of recovery, cells were transitioned to D10 supplemented with both doxycycline and puromycin to select for cells that have proper in-frame, full-length assemblies (GPR161). The GPCRome cell lines were also selected using puromycin. Following puromycin selection for two days, cells were transitioned to D10 and expanded before freezing down or used in subsequent assays.

### Fluorescence-activated cell sorting

For flow-based assays and cell sorting, frozen cell stocks were thawed and allowed to recover for several days in D10 medium. 48 hours prior to starting the experiment, cells were split into an appropriately sized dish such that they reach ∼75% confluency by the start of the sort. 36 hours prior to starting the assay, cells were induced with doxycycline hyclate (2 µg/mL). For GPR161 experiments, doxycycline was subsequently washed out after 24 hours, and cells were maintained in D10 for the remaining 12 hours prior to sorting. For GPCRome experiments, doxycycline was not washed out.

### GPR161 surface expression deep mutational scan

The surface expression sort assays were performed as follows. Following induction, cells were detached with TrypLE Express, washed, incubated with M2 FLAG APC-Surelight antibody (1:500, Abcam) for 1 h, washed 3x with PBS + 3% FBS, resuspended in BD FACS buffer, and kept covered on ice prior to sorting. Cell sorting was performed using a Cytoflex SRT. Briefly, cells were gated on FSC-A and SSC-A for HEK293T cells, then FSC-A and FSC-H for singlets. On the APC channel, gatesFor surface expression assays, the population was bimodal. Gates were set to divide the population into quartiles. For sorting experiments, we aimed to collect cells equal or greater than 100x the expected number of variants in our library.

### GPR161 cAMP signaling deep mutational scan

The cAMP assay was run as follows: cells were swapped to D10 (at indicated pH) with trimethoprim (10 μM) to stabilize the DHFR degron for 4 h. After this incubation, cells were detached using TrpyLE Express (Gibco), washed with PBS + 3% FBS, and resuspended in BD FACS buffer (BD). Cell sorting was performed using a Cytoflex SRT. Briefly, cells were gated on FSC-A and SSC-A for HEK293T cells, then FSC-A and FSC-H for singlets. For the cAMP assay, we assessed activity using eGFP on the FITC-A channel. Gates were set to divide the population into even quartiles, and we aimed to collect cells equal or greater than 100x the expected number of variants in our library.

### GPR161 whole-cell abundance deep mutational scan

The abundance assay was a modified VAMP-seq assay relying on complementation of mNeonGreen11 as a readout^42,43^. The GPR161 library was tagged with mNeon11. 48 h before sorting, a construct containing mNeonGreen-1-10 was transfected. This construct contains CMV-mNG2-1-10-P2A-mCherry, enabling selection and sorting of cells expressing this transiently transfected gene. A total of 18 wells (6 well plates) were transfected per assay using Lipofectamine 3000 following the manufacturer’s instructions. Induction of the library was performed using doxycycline, as for the surface expression and cAMP signaling screens. For sorting, cells were detached using TrpyLE Express (Gibco), washed with PBS + 3% FBS, and resuspended in BD FACS buffer (BD). Cell sorting was performed using a Cytoflex SRT. Briefly, cells were gated on FSC-A and SSC-A for HEK293T cells, then FSC-A and FSC-H for singlets. For the abundance assay, we first gated for mCherry+ cells and then assessed abundance using the mNeonGreen signal in the FITC-A channel. Gates were set to divide the population into even quartiles, and we aimed to collect cells equal or greater than 100x the expected number of variants in our library.

### GPCRome basal activity screen

The GPCRome library was plated into 10cm dishes such that after 48 hours, they reached ∼75% confluency. Upon plating, library expression was induced with doxycycline hyclate (2 µg/mL). After 24h, trimethoprim was added to the media (10 μM) to stabilize the DHFR degron for the remaining 24h prior to cell sorting. After this incubation, cells were detached using TrypLE Express (Gibco), washed with PBS + 3% FBS, and resuspended in PBS + 3% FBS. Cell sorting was performed using a BD FACSMelody. Briefly, cells were gated on FSC-A and SSC-A for HEK293T cells, then FSC-A and FSC-H for singlets. Next, BFP-cells were selected to gate for landing pad integrations. Finally, for the cAMP assay, we assessed activity using eGFP on the FITC-A channel. Gates were set to divide the population into even quartiles, and we aimed to collect cells with at least 100x the expected number of variants in our library.

### Genomic DNA extraction and sequencing

Following cell sorting, genomic DNA (gDNA) was extracted from cells using the Zymo Quick-DNA Miniprep Plus kit. For GPR161 mutational scanning experiments, all resulting gDNA was used as a template for PCR to amplify the target gene using the primers GGAGCAATTCCACAACACTTTTGTC and CGGCGAAAGCAGCTGCC. PCR reactions were then cleaned using Zymo DNA Clean and Concentrator-25 kits, mixed with NEB Purple Loading dye (6x, no SDS), and run on a 1% agarose 1xTBE gel. Target amplicons were excised and purified using Zymo Gel DNA Recovery kits. Amplicon DNA concentrations were then quantified using an Invitrogen Qubit dsDNA HS assay kit. Libraries were prepared for deep sequencing using the Illumina Nextera XT DNA Library prep kit. Libraries were indexed using the IDT for Nextera Unique Dual Indexes. Lengths of indexed libraries were quantified using the Agilent TapeStation HS D5000 assay, and concentrations were determined using the Invitrogen Qubit dsDNA HS assay kit. Samples were pooled and then paired-end sequenced (PE150) using either an Illumina NovoSeq 6000 or Novoseq X. For GPCRome screens, gDNA was extracted similarly. Receptor barcodes were amplified from the gDNA using the following primers: ATTGCATGAAGAATCTGCTTAGGGT and gcgggtttcttggatctatatgtggtc for 15 cycles. The resulting PCR reactions were cleaned using a Zymo DNA Clean and Concentrator-25 kit, and 10 ng of the resulting DNA was amplified using tcgtcggcagcgtcagatgtgtataagagacagATTGCATGAAGAATCTGCTTAGGGT and gtctcgtgggctcggagatgtgtataagagacagcggcatcgcgaagcttatCGCGTATATCTGGCCCGTACATC for 10 cycles to add Illumina Nextera adapter sequences. These reactions were cleaned up using the Zymo DNA Clean and Concentrator-5 kit. A final indexing PCR using Nextera Unique Dual Indexes was performed for 10 cycles. Lengths of indexed libraries were quantified using the Agilent TapeStation HS D5000 assay, and concentrations were determined using the Invitrogen Qubit dsDNA HS assay kit. Samples were pooled and then paired-end sequenced (PE150) using either an Illumina MiSeq i100.

### Next-generation sequencing data processing

Sequencing files were obtained from the sequencing core as fastq.gz after demultiplexing. The mutational scanning experiments were processed using a DMS-specific pipeline we developed, “Dumpling”, released as part of Rosace^49^. Dumpling implements the following steps: first, adapter sequences and contaminants were removed using BBDuk; then paired reads were error-corrected with BBMerge and mapped to the reference sequence using BBMap with 15-mers (all from BBTools ^50^). Variants in the mapped SAM file were called using the AnalyzeSaturationMutagenesis tool in GATK v4 ^51^. The output of this tool is a CSV containing the genotype of each distinct variant and the total number of reads for each sample. This was then further processed using a Python script, which filtered out sequences that were not part of the designed variants. The GPCRome fastq.gz files were processed using a custom pipeline (deposited on GitHub/Zenodo) to count receptor barcodes in each sample. We calculated receptor scores using a weighted mean-bin scoring system, then normalized them between 0 and 1.

### Mutational scan variant scoring

Variant scores were calculated with Lilace, a statistical model for scoring variants from FACS-based deep mutational scanning data ^44^. Lilace was run with default settings, in which within-screen synonymous variants are used as the baseline. Scores were centered at zero for synonymous variants; gain-of-function variants (increased fluorescent signal relative to synonymous) were positive, and loss-of-function variants (decreased signal) were negative. Throughout this work, we use a cutoff of two standard deviations of the synonymous variant mean to define loss-of-function, but we note there are more sophisticated methods, such as COSMOS^52^, which may further resolve phenotype-specific variant effects.

## Supplemental figures

**Supplemental Figure 1:**
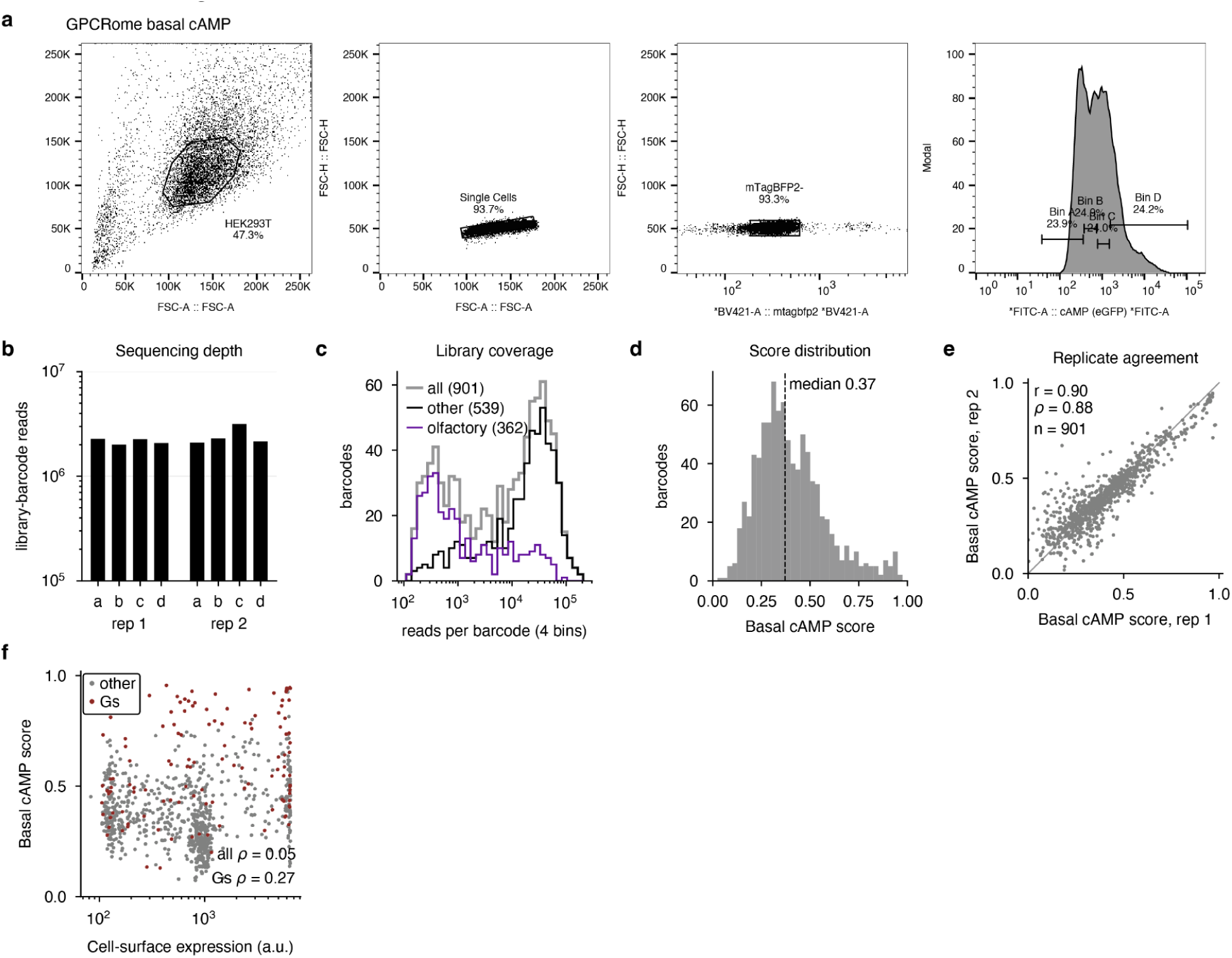
GPCRome sort experiments. (a) Example FACS gating strategies for the GPCRome basal cAMP signaling screen. (b) Total number of receptor barcode reads observed per sort bin across each replicate. (c) Total reads per barcode. We note that, similar to previous work, olfactory receptor barcodes are depleted relative to other GPCRs^9^. (d) Basal cAMP score distribution across the library. (e) Replicate correlation plot between replicates. (f) Comparison of cell surface expression from Tedman et al. 2026^9^ and basal cAMP signaling scores (this study).

**Supplemental Figure 2:**
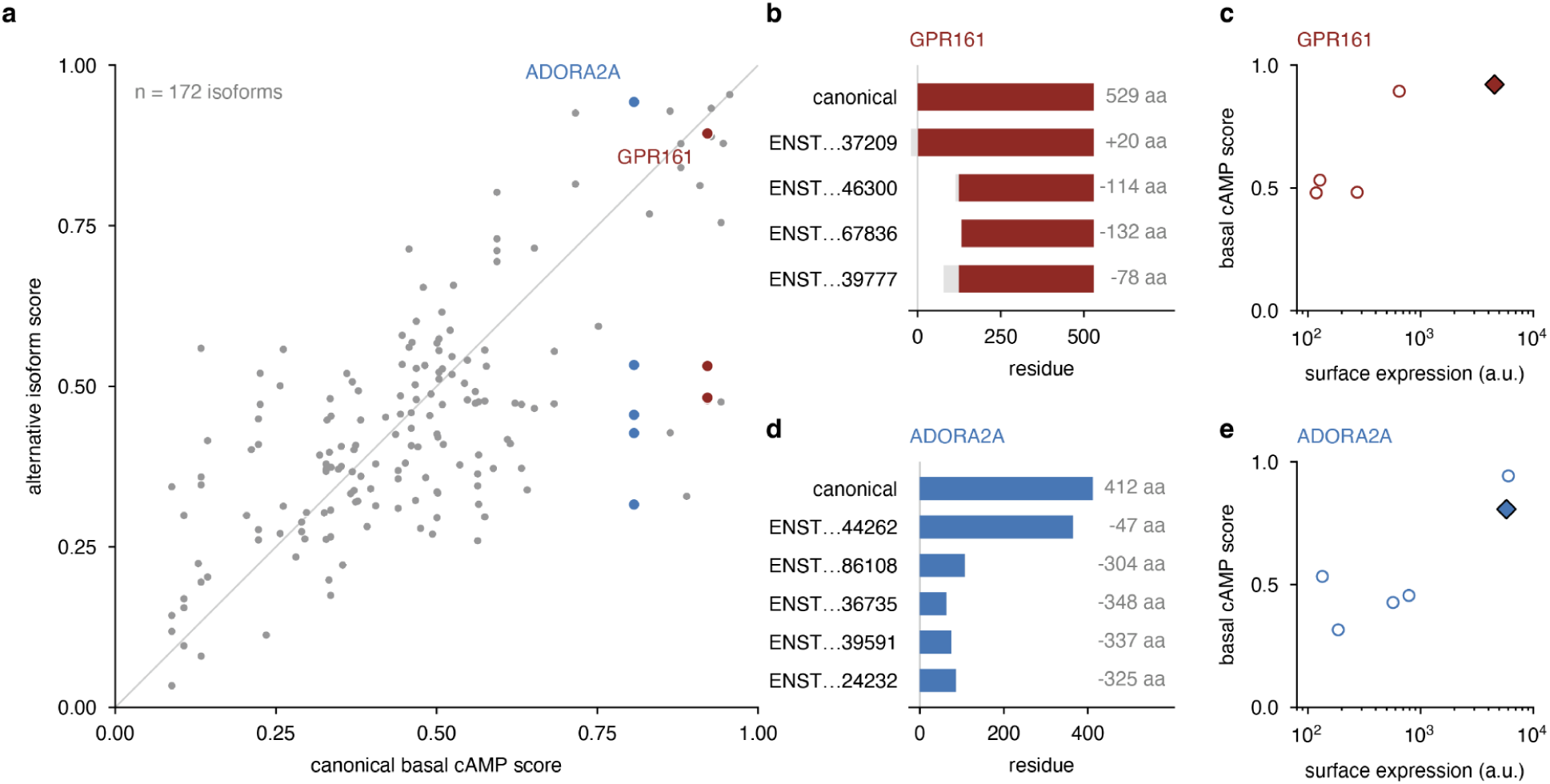
GPCR isoforms with altered basal cAMP signaling. (a) Scatter plot comparing basal cAMP scores between canonical receptors and their isoforms. (b) Sequence map of GPR161 and its four isoforms. (c) Scatter plot comparing basal cAMP signaling and cell surface expression for GPR161 isoforms. (d) Sequence map of ADORA2A and its four isoforms. (e) Scatter plot comparing basal cAMP signaling and cell surface expression for ADORA2A isoforms.

**Supplemental Figure 3:**
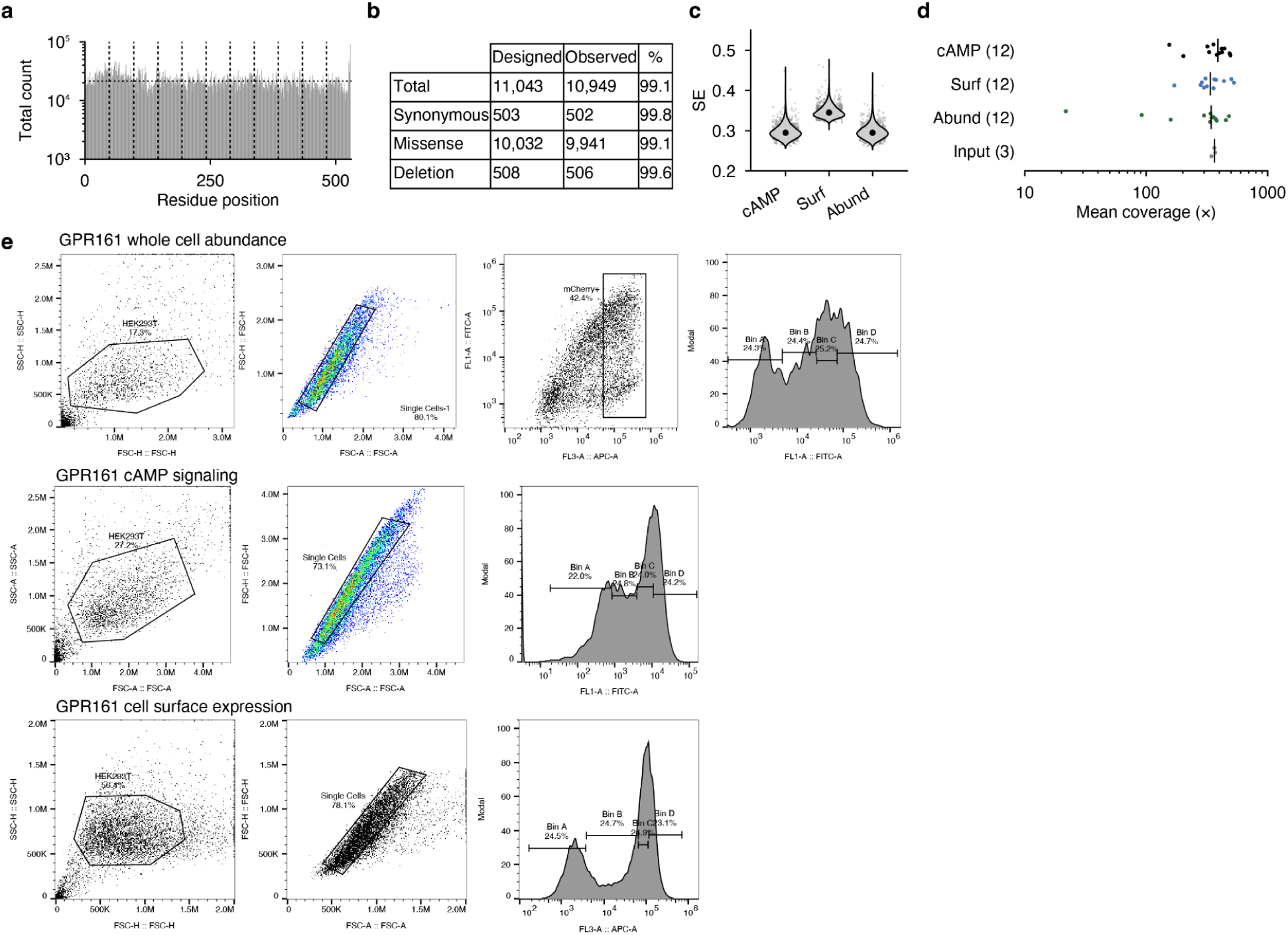
GPR161 mutational scanning sort experiments and quality control. (a) Positional coverage of the GPR161 mutational library (b) Table of designed and observed variants in the GPR161 mutational library (c) SE violin plots for each GPR161 mutational scanning experiment. (d) Average count per variant in each condition screened, stratified by experiment type. Each point corresponds to a bin from each sorting experiment. (e) Example FACS gating strategies for the GPR161 abundance, cAMP signaling, and surface expression screens.

## Supplemental tables

**Supplemental Table 1: GPCRome basal cAMP scores**

Each row is one barcoded construct. basal_camp_score is the weighted mean sorting score for each receptor; basal_camp_score_sd is the SD between them and rep1_score/rep2_score the per-replicate values. n_replicates is how many sorts passed the ≥50-read threshold. canonical_rank ranks the 727 canonical constructs by score. coupling, gs_primary and gs_any are IUPHAR/Guide to PHARMACOLOGY transducer annotations. surface_expression_tedman is plasma-membrane expression from Tedman et al. 2026^9^; surface_value_saturated flags values ≥4000, where that estimate saturates and is a lower bound. prior_constitutive_evidence and the prior_evidence_* columns are hand-curated; only the top 40 canonical constructs were reviewed.

**Supplemental Table 2: Oligonucleotide sequences used in GPR161 library construction**

Each row is one oligonucleotide with a designed variant or a primer used to amplify variant subpools or the WT backbone of the GPR161 gene as part of the DIMPLE protocol.

**Supplemental Table 3: GPR161 deep mutational scanning scoreset**

Variant effect scores from the GPR161 deep mutational scan on protein abundance, surface expression, and cAMP signaling.

